# Block-structured Bayesian source estimation model for magnetoencephalography signals

**DOI:** 10.1101/2025.05.30.656951

**Authors:** Kai Miyazaki, Okito Yamashita, Yoichi Miyawaki

**Affiliations:** Graduate School of Informatics and Engineering, The University of Electro-Communications, Tokyo, Japan; Advanced Telecommunications Research Institutes International, Kyoto, Japan; RIKEN Center for Advanced Intelligence Project, RIKEN, Tokyo, Japan; Center for Neuroscience and Biomedical Engineering (CNBE), The University of Electro-Communications, Tokyo, Japan

**Keywords:** Magnetoencephalography, Inverse problem, Hierarchical Bayesian model, Block-structured covariance

## Abstract

The human brain consists of functionally specialized areas that process specific types of information and interact with one another. Magnetoencephalography (MEG) is a neuroimaging technique that captures brain activity at high temporal resolution. However, its spatial resolution is insufficient to accurately localize neural activation, and thus cannot capture the interactive nature of human brain activity. To resolve this issue, various MEG source estimation models have been proposed. In particular, models incorporating prior information from functional magnetic resonance imaging (fMRI), which offers superior spatial resolution, have improved source estimation accuracy. However, these models typically ignore the similarity of activity across different brain areas, and still fall short in tracking dynamic brain activity pattern at a sufficient spatial scale. In this study, we developed a block-structured model that integrates information about functional areas and the inter-areal relationships of the human brain into MEG source estimation based on a hierarchical Bayesian framework. We evaluated our model performance using simulation data with sequential activation across multiple brain areas. Results showed that our model outperformed conventional approaches in source estimation accuracy, suggesting that incorporating the functional areal information and inter-areal relationships may enhance MEG source estimation, enabling human neuroimaging at high spatiotemporal resolution.

## 1 Introduction

The human brain processes external information at fine spatial and temporal scales through dynamic and harmonious interaction among multiple brain areas. Neuroimaging techniques require a sufficient spatiotemporal resolution for tracking such neural processes. Functional magnetic resonance imaging (fMRI) and magnetoencephalography (MEG) are widely used neuroimaging techniques for measuring brain activity. fMRI offers high spatial resolution, capable of even distinguishing cortical layer structures at the submillimeter level. However, its temporal resolution is limited due to its reliance on blood oxygenation changes. In contrast, MEG provides high temporal resolution by detecting magnetic fields generated by electrical neural activity, but its spatial resolution is constrained because the sensors are placed above the scalp and detect signals from multiple brain areas. Consequently, there are no neuroimaging techniques that can sufficiently track neural activity with both high spatial and temporal resolution.

To resolve this issue, previous studies have attempted to estimate the locations of neural sources in the brain that generate the measured MEG signals [1–3]. This approach is known as MEG source estimation, and various source estimation models have been developed and evaluated. The most widely used method moderates the ill-posedness of the inverse problem by introducing regularization techniques, as the number of MEG sensors is much smaller than the number of candidate source localizations. Minimum norm estimation (MNE) [1] employs squared regularization, resulting in stable but spatially diffuse solutions. In contrast, automatic relevance determination (ARD) [4, 5] imposes a sparse prior within Bayesian framework to estimate more focal sources. In addition, high spatial resolution information from fMRI can be incorporated to improve MEG source estimation. For example, fMRI-derived blood oxygenation contrasts can serve as prior information. Variational Bayesian multimodal encephalography (VBMEG) [3] incorporates such information as a hierarchical prior within a Bayesian framework. Although these models have achieved a certain level of accuracy in source localization, their estimation processes are limited in that they do not consider the functional organization of the human brain [6–8]. Specifically, they treat each cortical location independently, without incorporating the spatial similarity of neural activity within the same functional areas or interaction among multiple brain areas [9–12].

To address this limitation, we hypothesized that incorporating such structural and functional properties into the prior distribution for MEG source estimation would lead to improve its accuracy. Specifically, we incorporated information of functional area and inter-areal relationships into the model as a hierarchical prior within a variational Bayesian framework. The model is expected to produce source estimates that more faithfully reflect the underlying functional organization of the brain, thereby enabling accurate MEG source estimation.

## 2 Method

### 2.1 MEG source estimation

MEG signals were computed using a forward model based on the positional relationship between the signal source and the MEG sensors. The forward model is described as

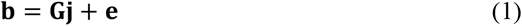

where **b** (ℝ^*N*×*T*^ matrix; *N*, number of MEG sensors; *T*,number of time points) denotes the measured MEG signal, **G** (ℝ^*N*×*M*^ matrix; *M*, number of vertices on the cortical surface) is the leadfield matrix that describes how each cortical source current contributes to the MEG signal at each sensor location, **j** (ℝ^*M*×*T*^ matrix) represents the cortical source current, and **e** (ℝ^*N*×*T*^ vector) is the sensor noise modeled as a zero-mean multivariate Gaussian random variable with covariance *β*^−1^**Σ**_***ϵ***_ . **G** was computed using VBMEG software.

MEG source estimation is to estimate **j** given **b** and **G** in Eq. (1). However, because the number of vertices typically exceeds the number of MEG sensors, it is an ill-posed problem and a unique solution cannot be obtained. To resolve this issue, various models have been proposed. In this paper, we developed a Bayesian estimation model that incorporates group information based on functional areas in the brain and inter-areal relationships as a hierarchical prior distribution. To evaluate whether these properties improve source estimation accuracy, we compared four models: (1) a hierarchical Bayesian model assuming that cortical surface vertices are treated independently of each other (“hierarchical automatic relevance determination” model), (2) a hierarchical Bayesian model incorporating only functional areal grouping (i.e., grouping vertices according to cortical parcellation; “hierarchical grouped automatic relevance determination” model), (3) a hierarchical Bayesian model incorporating both functional areal grouping and inter-areal relationships (i.e., grouping vertices according to cortical parcellation and defining interactions between groups; “block-structured hierarchical Bayesian estimation” model), and (4) a model with L2-norm regularization (minimum norm estimation, MNE) as a widely used standard method.

#### Hierarchical automatic relevance determination

The Bayesian estimation model introduces both the likelihood and the prior distribution for **j** as Gaussian distributions:

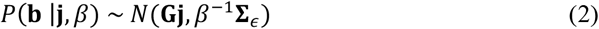

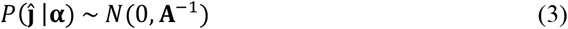

where *β*^−1^**Σ**_***ϵ***_ represents the sensor noise variance, and **A** is the ℝ^*M*×*M*^ source covariance matrix. If we put a uniform prior on the diagonal elements *α* of the source covariance **A**, we will obtain automatic relevance determination (ARD) [4, 5]. To improve the source estimation accuracy, a hierarchical prior can be placed on the elements of *α*. We call this model hierarchical automatic relevance determination (hARD). hARD places a conjugate prior distribution on the elements *α* in the form of the following Gamma distribution:

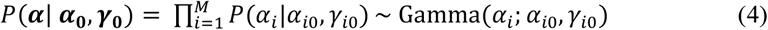

where *α*_*i*0_ represents the relevance parameter of each vertex, and the *γ*_*i*0_ represents the degree of freedom. Based on these probability distributions, the posterior probability distribution is then defined as:

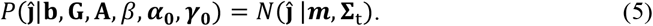

To estimate approximate posterior distribution, we used the variational Bayesian estimation and mean field approximation [13, 14]. The mean ***m*** and covariance **Σ**_***t***_ are given by

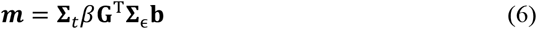

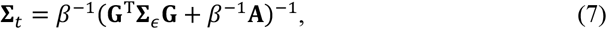

respectively. The estimated current ĵis obtained by calculating the expectation of this posterior distribution. hARD optimizes the hyperparameter as the following update rules:

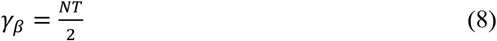

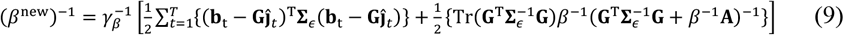

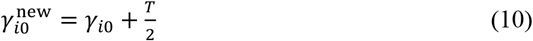

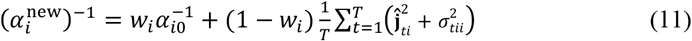

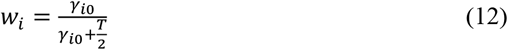

where 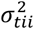 is diagonal element of **Σ**_*t*_ in Eq. (7), and *w*_*i*_ is the prior weight, which controls the balance between the first and second terms of Eq. (11) based on the confidence of hyperparameter of hierarchical prior (in the case of ARD, *w*_*i*_ is 0). By iteratively applying the update rules shown in Eqs. (6–12) until convergence, the model parameters are optimized and an approximate posterior distribution of the cortical source current is obtained. Figure 1 (a) shows the graphical illustration of hARD.

**Fig. 1.**
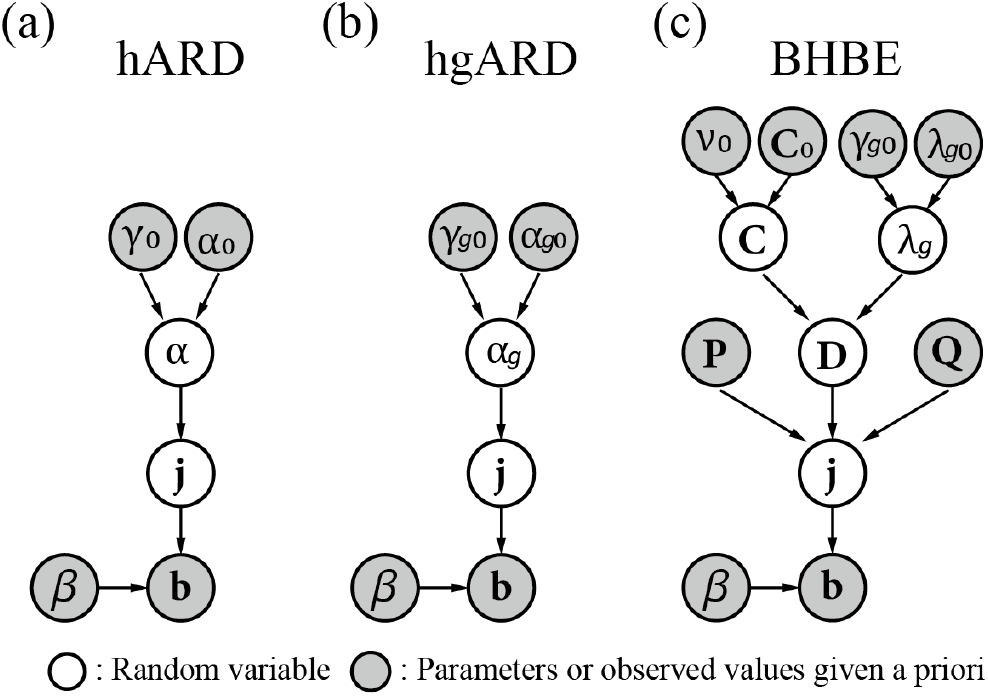
Graphical representations of the Bayesian models.

#### Hierarchical grouped automatic relevance determination

hARD treats each vertex independently and ignores the fact that vertices within the same functional area of the brain may exhibit similar activation due to shared processing roles. To incorporate this principle of functional organization into the model, we developed a Bayesian estimation model that introduces grouping of vertices based on cortical parcellation. We call this model hierarchical grouped automatic relevance determination (hgARD). The likelihood and prior distribution for **j** is the same as Eqs. (1-2), and the diagonal elements *α* of the source covariance matrix are shared within the group as

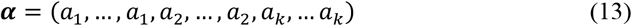

where *k* represents the number of groups. The number of repetitions of each *a*_*k*_ in *α* corresponds to the number of vertices assigned to the *k*-th group. The conjugate prior distribution for this grouped element *α* is given by the following Gamma distribution:

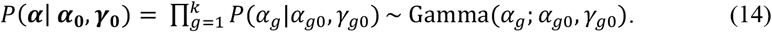

Based on these probability distributions, the posterior probability distribution and the mean ***m*** and covariance **Σ**_***t***_ are same as Eqs. (5-7). hgARD optimizes the hyperparameters using an update rule as in hARD. The update rules are similar to Eq. (10-12) but are modified according to Eq. (13). In update rules of Eq. (10-12), *γ*_*i*0_, *α*_*i*0_ and *T* are replaced by *γ*_*g*0_ *α*_*g*0_ and *N*_*g*_*T*, respectively, where *N*_*g*_ is the number of vertices belonging to the *g*-th group. Moreover, the term 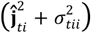 is replaced by the group-wise sum 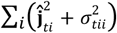 By iteratively applying the update rules until convergence, the model parameters are optimized and an approximate posterior distribution of the cortical source current is obtained. Figure 1 (b) shows the graphical illustration of hgARD.

#### Block-Structured Hierarchical Bayesian estimation model

hARD and hgARD treat only the diagonal elements of the source covariance matrix that represent source current variance at each cortical location, and ignore the off-diagonal elements that represent relationships between functional areas. To consider a full source covariance matrix, the conjugate prior distribution should be Wishart distribution [5]. Although a full covariance matrix can potentially account for the covariance structure of the source currents, it is impractical to estimate due to the excessive number of parameters. To mitigate this difficulty, we developed a Bayesian estimation model that incorporates grouping not only for the diagonal elements but also for the off-diagonal elements of the source covariance matrix. We call this model Block-Structured Hierarchical Bayesian estimation (BHBE) model, which corresponds to the model (3). The likelihood and prior distribution for **j** is the same as Eqs. (1-2), and the source covariance matrix **A** is defined as *k* × *k* block matrices

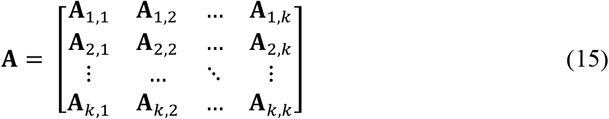

where the diagonal blocks **A**_*l,l*_ and the off-diagonal blocks **A**_**l**,**l**_*′* are defined as

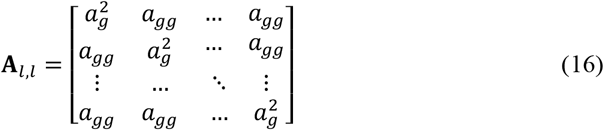

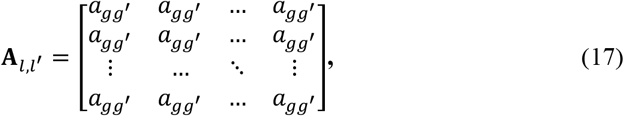

respectively (*g* ≠ *g′* and *g*,*g′* = 1,2, …, *k*). According to the method proposed by Archakov and Hansen [15], the block-structured covariance matrix **A** can be decomposed as

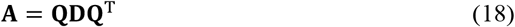

where **Q** is an orthonormal matrix and **D** is a ℝ^*M*×*M*^ block-diagonal matrix. **Q** is defined as

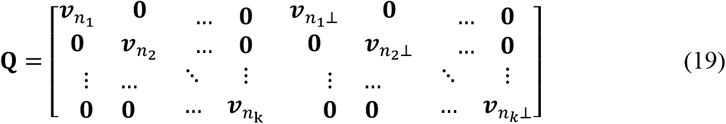

where 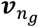 is 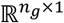 column vector with entries 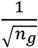 and 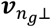 is 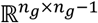 matrix that is orthogonal to 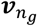 such that 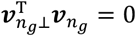 and 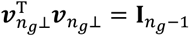. **D** is defined as

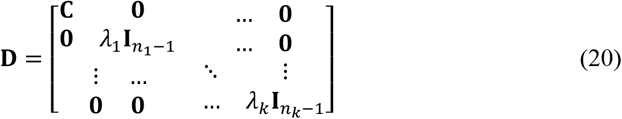

where **C** is a ℝ^*k*×*k*^ positive definite matrix representing the relationship between the groups, and *λ*_*g*_ is a positive scalar. The elements of the original covariance matrix **A** can then be expressed as

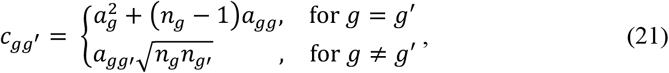

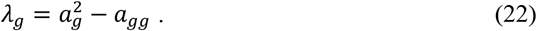

Here the diagonal blocks of **A** are determined by the diagonal elements of **C** and *λ*_*g*_ and the off-diagonal blocks of **A** are controlled by the off-diagonal elements of **C**. From Eq. (22), *λ*_*g*_ represents the difference between the diagonal and off-diagonal elements within group *g*, reflecting how much the variances 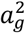 of the diagonal elements deviate from covariance *a*_*gg*_ of the off-diagonal elements within the group. This indicates that as *λ*_*g*_ increases, correlations among vertices within the group decreased, resulting in greater variability in the estimated currents among those vertices. In the current definition of **A** in Eq. (15), the elements are arranged in group-wise order (e.g., diag(**A**) = (*a*_11_, …, *a*_11_, *a*_22_, …, *a*_22_ … *a*_*kk*_, …, *a*_*kk*_)). However, in actual data, the vertex grouping doesn’t necessarily follow such an ordered structure, and the elements of **A** may appear in irregular arrangement (e.g., diag(**A**) = (*a*_11_, *a*_22_, *a*_*kk*_, *a*_11_, *a*_*kk*_, … *a*_11_)). To align the group ordering to the vertex ordering, we introduced the permutation matrix **P**, and defined the prior distribution as

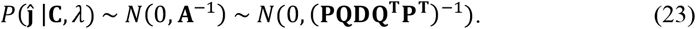

The conjugate prior distributions for this hyperparameter **C** and *λ* are given by the following Wishart and Gamma distributions:

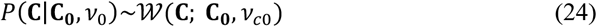

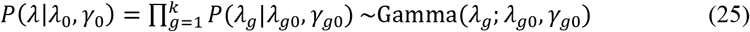

where *λ*_*g*0_ is a hyperparameter that represents the expected difference between the variance and covariance within the group, the *γ*_*g*0_ is the degree of freedom and the **C**_0_ ∈ ℝ^*k*×*k*^ defines the covariance between groups. Based on these probability distributions, the posterior mean ***m*** and covariance **Σ**_***t***_ are given by Eqs. (8,9), respectively. Whereas the previous study [16] used Markov-chain Monte Carlo algorithm to explore the posterior distribution, we employed the variational Bayesian estimation and the mean field approximation. The hyperparameters *λ* and **C** are then optimized by the following update rules

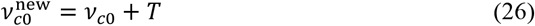

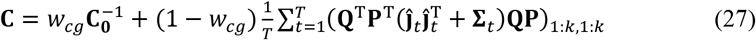

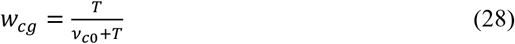

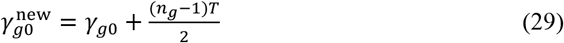

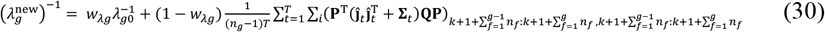

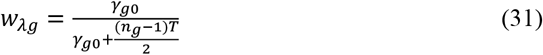

where (・) _1:*n*,1:*n*_ denotes the submatrix formed by selecting rows and columns from 1 to *n*. In this formulation, the number of blocks determines the number of the groups (cortical parcellation), and the hyperparameter **C**_0_ and *λ*_*g*0_ determine the relationship between and within the groups, respectively. Figure 1 (c) shows the graphical illustration of the proposed model.

### 2.2 Experiments

We performed MRI and MEG experiments to obtain the cortical surface model (from MRI data) and the head position in MEG scanner (from MEG data) to be used in the simulation analysis according to the following procedures.

#### Participant

One individual took part in both MRI and MEG experiments. The participant gave written informed consent before the experiment. The procedure was approved by the institutional review board of the University of Electro-Communications, Advanced Telecommunications Research Institute International (ATR) Brain Activity Imaging Center, and Ethical Review Board of Osaka University Hospital. The experiment was conducted under corrected vision with MRI-compatible glasses.

#### MRI experiment

Anatomical images were obtained using an MRI system (3-Tesla MAGNETOM Prismafit, Siemens) located at the ATR Brain Activity Imaging Center. The magnetization-prepared rapid-acquisition gradient-echo (MP-RAGE) sequence (208 sagittal slices; TR, 2,250 ms; TE, 3.06 ms; FA, 9 deg; FOV, 256 × 256 mm; voxel size, 1.0 × 1.0 × 1.0 mm) was used.

#### MEG experiment

The participant’s head and the positions of the five marker coils (three on the forehead and two on each earlobe) were first recorded using a 3D scanner and a position sensor system (FastSCAN Cobra, Polhemus Inc., USA). These markers were used to calibrate the head position relative to the MEG sensors inside the MEG scanner (PQ1160C, Yokogawa Electric Co., Japan). Then, the participant was asked to lie on their back in the MEG scanner and remain as still as possible. The current was then applied to the five marker coils to generate a magnetic field, and the magnetic field was measured with 160-channel superconducting quantum interference device (SQUID) sensors in the MEG scanner. The spatial relationship between the MEG sensors and the marker coils was computed by solving an inverse problem based on the measured magnetic field. Using the position information of the marker coils, the participant head location relative to the MEG sensor locations was estimated. Finally, the high-resolution anatomical MRI images of the participant were aligned to the participant head locations in the MEG scanner.

#### Cortical parcellation for grouping

We employed a standard brain atlas provided by the Human Connectome Project (HCP) dataset [7], which divides the cortical surface into 180 areas per hemisphere, resulting in 360 areas across both hemispheres, each of which serves as a group information for MEG source estimation in this study. To assign these areas for the participant cortical surface, we used the Freesurfer software (http://surfer.nmr.mgh.harvard.edu/) to normalize the participant’s MRI structure data to the standard brain space and the cortical surface was extracted from the normalized participant brain using the polygon model with 15,002 vertices at the gray/white matter outer boundary. Then each vertex was assigned with a label for the corresponding area. There were exceptional vertices that did not correspond to any of the HCP-defined areas, which were located near the corpus callosum. These vertices were labeled with the 361st area (1,110 vertices were identified in total).

### 2.3 Simulation data generation and the hyperparameter

#### Cortical source current model

To evaluate source estimation accuracy, we simulated temporally localized and spatially distinct cortical activations across early visual areas. In this simulation, three regions of interest (ROIs) were defined: V1, V2, and V3, and they were sequentially activated as shown in the left panel of Fig. 2. Each ROI’s cortical source current followed a sinusoidal waveform with a half-cycle duration of 25 ms, peaking at 13 ms (V1), 25 ms (V2), and 37 ms (V3), respectively. This waveform was assigned to each ROI in left or right hemisphere and its polarity was reversed in the opposite hemisphere (the positive activation was assigned to the left hemisphere in condition 1 and the polarity was reversed in condition 2).

**Fig. 2.**
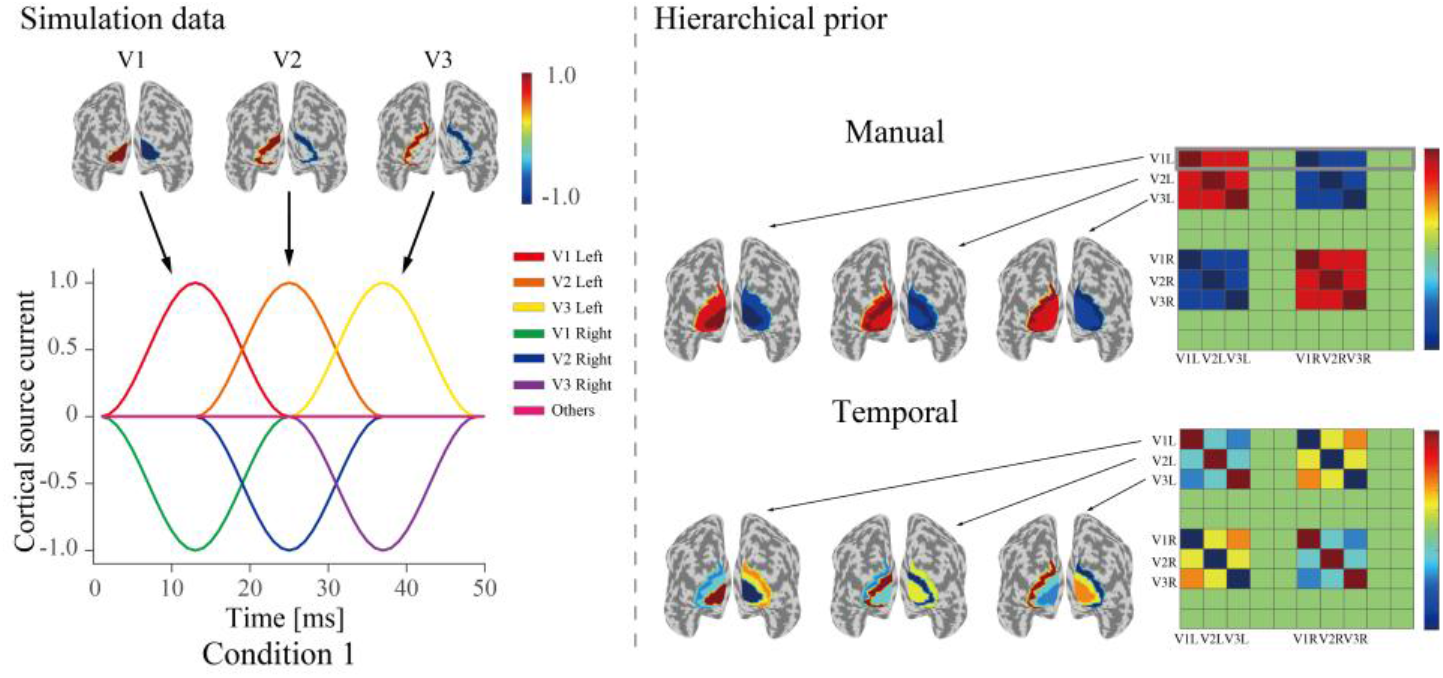
Schematic view of cortical source current model and hierarchical prior.

#### Sensor noise model

The noise was independently added to each MEG sensor, each time point, and each trial. It was sampled from a Gaussian distribution with a mean of 0 and standard deviation σ. The value of σ was adjusted such that the resulting signal-to-noise ratio (SNR) varied from −5 dB to 5 dB with 5 dB steps. The SNR was calculated using the following equation:

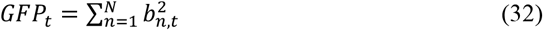

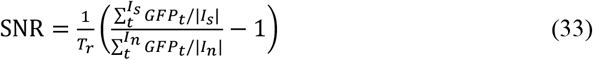

where *GFP*_*t*_ represents the global field power at time *t, b*_*n,t*_ denotes the MEG signal recorded from the *n*th sensor at time *t, T*_*r*_ denotes the number of trials, and *I*_*s*_ and *I*_*n*_correspond to the task and noise time windows, defined as periods during which the brain was assumed to be active and inactive, respectively. The resulting SNR was then converted to decibels (dB) using 10 log_10_(SNR). To compute the SNR, the task time window was defined as 1-25 ms, while the noise time window was defined as 26–50 ms using only the added noise components. One simulation dataset in this scenario consisted of five runs, each containing ten trials per condition, resulting in a total of 100 trials (50 trials per condition).

#### Hyperparameters for hierarchical priors

To define the hyperparameters for hierarchical priors, we used two types of covariance matrices (Fig. 2, right): (1) a manually designed covariance matrix, and (2) a temporal covariance matrix. For the manually designed covariance matrix, the diagonal elements corresponding to the active ROIs (V1, V2, and V3) were set to 1, and all others to 0. The off-diagonal elements were set as follows: 0.8 for ROI pairs within the same hemisphere (e.g., left V1 and left V2), – 1 for ROI pairs in opposite hemispheres corresponding to the same functional area (e.g., left V1 and right V1), and –0.8 for ROI pairs in opposite hemispheres corresponding to different functional areas (e.g., left V1 and right V2). The temporal covariance matrix was computed by calculating the covariance across time from the simulated cortical source current. These two covariance matrices were used as hyperparameters **C**_0_ in the hierarchical Bayesian models. For hARD and hgARD that omit to model inter-areal relationships, only the diagonal elements of these matrices were retained. In this simulation, the ideal value of *λ*_*g*_ is zero because the source current is uniform within each ROI. However, directly setting it to zero would cause a division by zero during the estimation process. Therefore, we set the value of *λ*_*g*_ to 1.0 × 10^−3^ in this study. To examine the effect of the hierarchical prior in estimation, the prior weight was varied from 0 to 0.05 with 0.01 steps.

#### Quantitative evaluation of models

To evaluate the estimation accuracy, we computed the area under the precision-recall curve (APR) [17] at the peak activation time point of each ROI, using the absolute values of the estimated currents normalized by the maximum across all vertices at the corresponding time point. In addition, we computed the spatiotemporal correlation between the estimated current and the cortical source current across all vertices and time points to assess the overall agreement. Before computing each metric, the estimated source currents were averaged across all 50 trials for each condition (i.e., condition 1 and condition 2).

## 3 Results

### 3.1 Simulation data analysis

Figure 3 shows the spatial distribution of the estimated current. The color represents estimated current amplitudes at the time points indicated at the top. The color scale was symmetrically adjusted based on the absolute maximum of the estimated current amplitude. Values between 0 ±10% of the absolute maximum of the estimated current amplitude were not displayed on the map, to emphasize the most salient regions of activation. Representative results at −5 dB SNR under condition 1 using a prior weight of 0.03 for the two types of hierarchical priors are shown in this figure. The results demonstrate that BHBE accurately estimated the current distribution, closely matching the true source in both spatial location and polarity. In contrast, the other models produced scattered estimates with frequent polarity reversals in neighboring vertices.

**Fig. 3.**
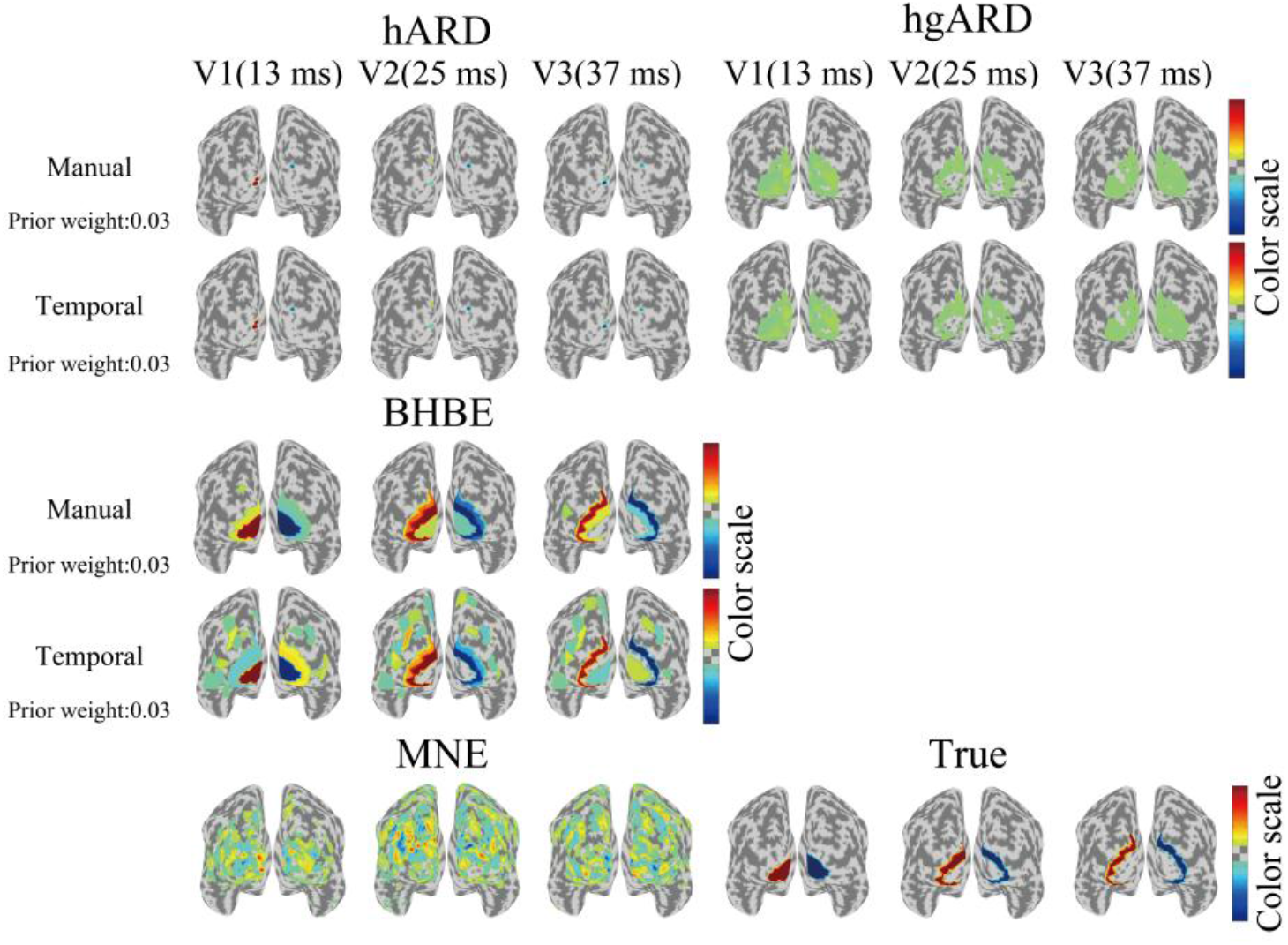
Estimated cortical current maps.

Figure 4 shows the waveforms of the estimated current. The results show that BHBE accurately reconstructed the three distinct waveforms corresponding to V1, V2 and V3. In contrast, the other models produced noisy estimates, with a slight increase in amplitudes around the activation period of V1 (1–25 ms), but failed to reconstruct waveforms corresponding to the activation periods of V2 and V3.

**Fig. 4.**
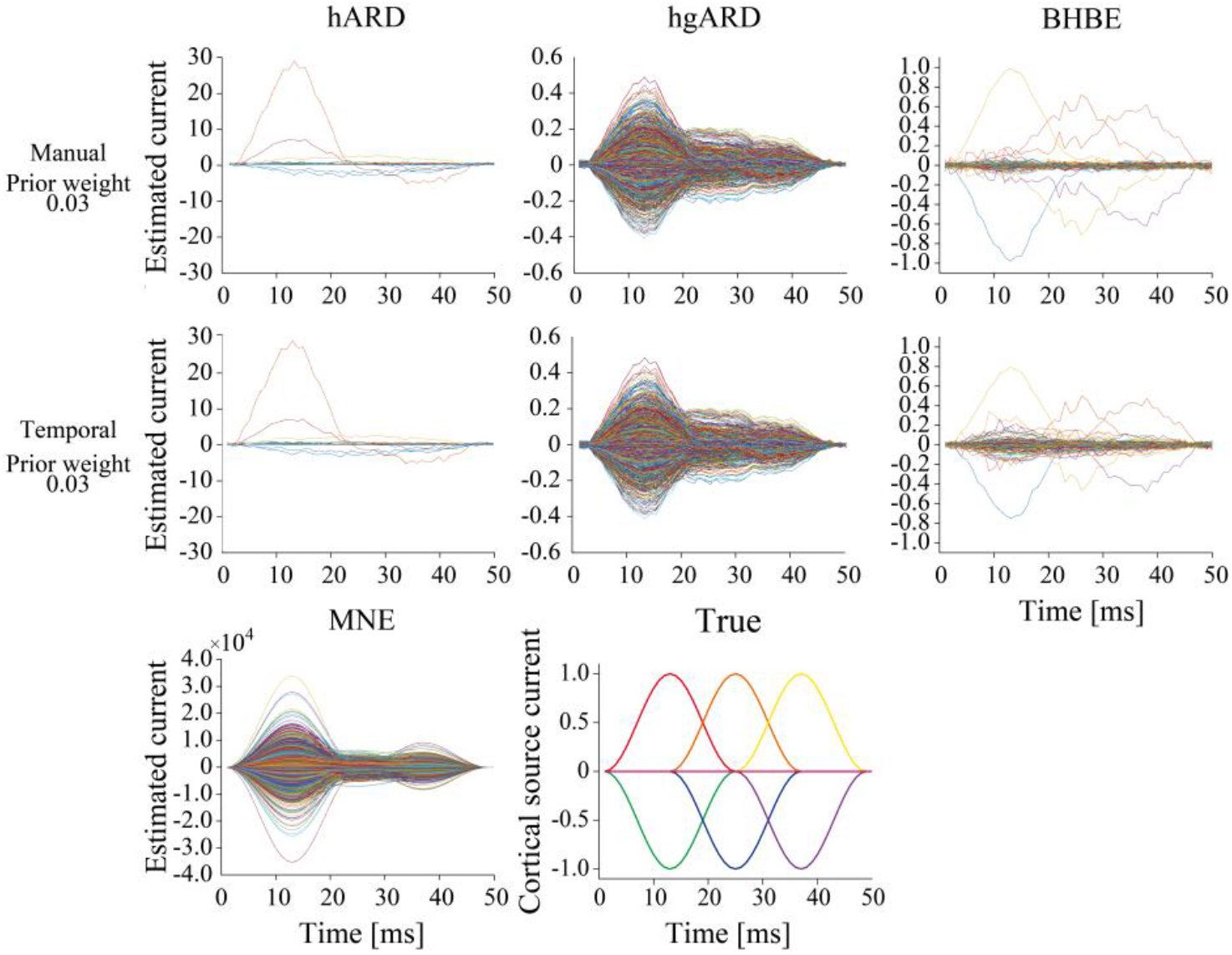
Waveforms of estimated cortical current.

Figure 5 shows the APR plotted against the prior weight (results from condition 1 are shown as representative examples). Since MNE is not a hierarchical Bayesian model, a single APR value was obtained and plotted across all prior weight for comparison. The results show that BHBE achieved higher APR values for all tested ROIs, indicating that estimated sources closely match the true source distribution, than all other models.

**Fig. 5.**
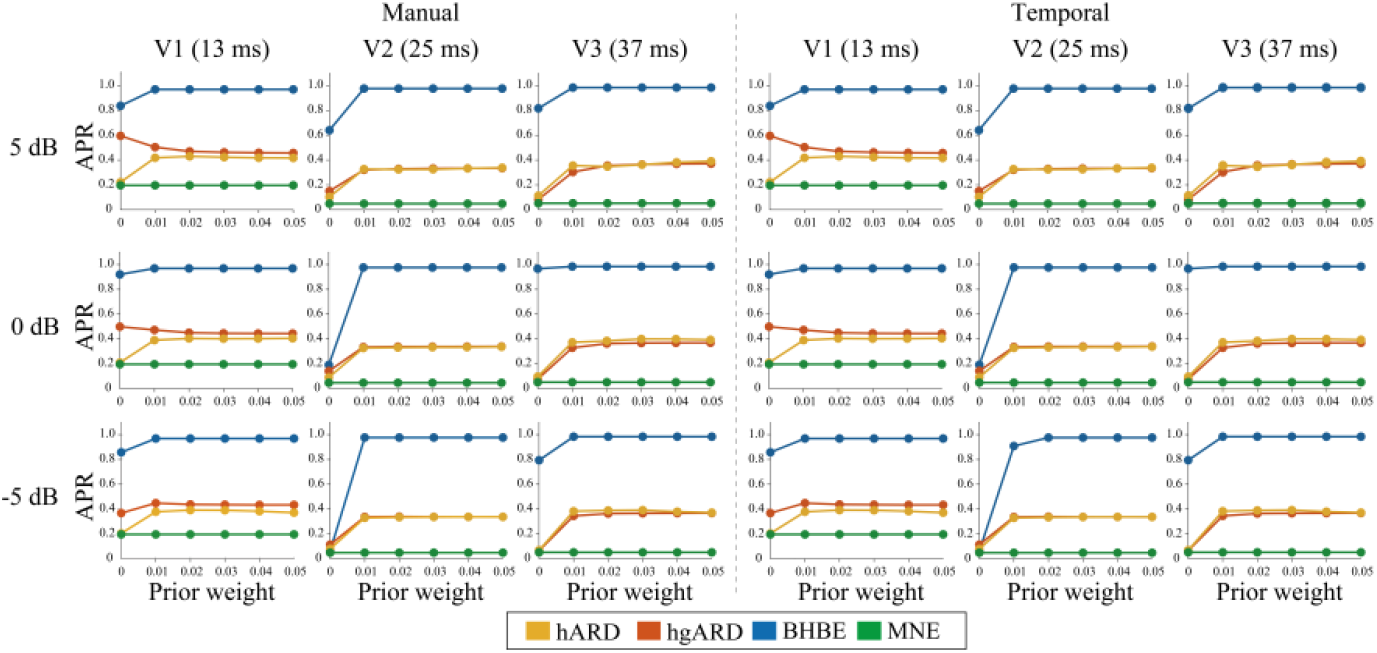
APR of the estimated cortical current at the peak time point of each ROI.

Figure 6 shows spatiotemporal correlation between the true and estimated cortical currents, plotted against the prior weight. The results demonstrate that BHBE outperformed the other models irrespective of the prior weight, indicating that our proposed model can accurately estimate the source cortical current with high fidelity in both space and time.

**Fig. 6.**
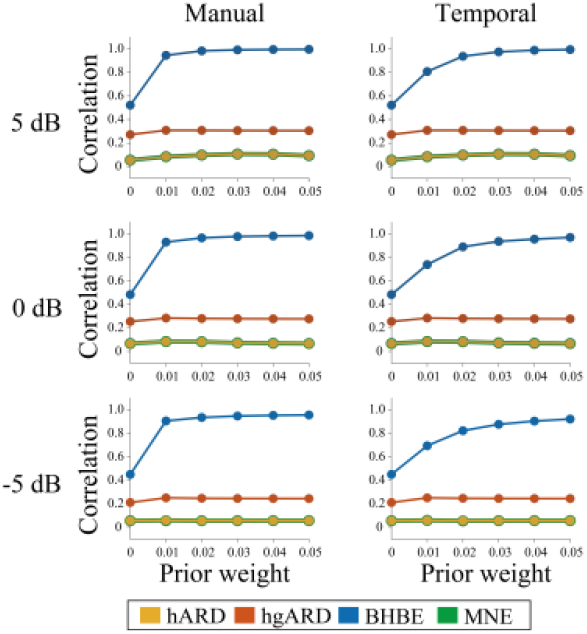
Spatiotemporal correlation between the true and the estimated source cortical current.

## 4 Conclusion and Discussion

In this study, we proposed a block-structured hierarchical Bayesian estimation (BHBE) model to improve the source estimation accuracy by incorporating both functional areal grouping and inter-areal relationships. We compared it with models that incorporated only functional areal grouping and neither of them. According to the simulation results, BHBE outperformed the other models in source estimation accuracy. These findings suggest that incorporating functional areal grouping and inter-areal relationships into MEG source estimation models can significantly enhance the accuracy of cortical source current estimation. Although the formulation is different from this study, a block-structured model has also been explored in a previous study [16]. A key difference lies in the type of information incorporated: whereas the previous study used physical relationships by grouping spatially adjacent vertices into blocks, our model incorporates functional relationships by grouping vertices according to a widely accepted cortical parcellation derived from fMRI data [7]. As a result, our approach may be able to capture inter-areal dependencies that are not necessarily spatially adjacent but are functionally connected.

However, there might be a possibility that the present simulation setups were suitable for BHBE, potentially biasing the results of source estimation. In this study, the cortical source current was defined such that all vertices within each ROI exhibited uniform activation with the same amplitude, which may favor our model’s framework of assigning the shared hyperparameters to vertices within the same group. To further test whether our model can generalize to various situations, it would be necessary to evaluate our model with more complex patterns of source cortical current. Although such complex activation patterns were not examined in this study, the flexible prior structure of our model may help address such situations. Theoretically, increasing the value of *λ*_*g*_ reduces the correlation of the estimated current within the same group, thereby increasing the variance of the estimated currents within the group. Thus, by adjusting *λ*_*g*_ our model is expected to flexibly accommodate a range of activation patterns, from homogeneous to more complex heterogeneous ones.

Although we used either a manually designed covariance matrix or a temporal covariance matrix as the hierarchical prior in this study, other sources of information may also be valuable for real data analysis. For example, fMRI experiments could be performed with the same experiment design as MEG, allowing the spatial and temporal covariance structures of the fMRI signals to be used as hierarchical prior distributions. By incorporating such neuroscientifically grounded priors to define inter-areal relationships, we might be able to analyze when and where each brain area is active and interacts during the processing of sensory information. Thus, our model framework may offer new opportunities for capturing dynamics of human neural activity with high spatiotemporal resolution.

## Acknowledgments

This study was partially funded by Research Fellow of Japan Society for the Promotion of Science (JSPS) Grant Number 24KJ1127, JSPS KAKENHI Grant Number JP18KK0311, JP20H00600, JP25H01138, a research granted from Murata Science and Education Foundation, the Naito Foundation, and Moonshot Program 9 under Grant JPMJMS2291.

